# Oral immunization of broilers with chitosan nano-encapsulated extracellular and cell wall proteins of necrotic enteritis-induced *Clostridium perfringens*

**DOI:** 10.1101/2020.10.23.351817

**Authors:** Nour Muinis Ramadan, Gabriel Akerele, Sankar Renu, Gourapura Renukaradhya, Ramesh Selvaraj

**Affiliations:** Department of Poultry Science, College of Agricultural and Environmental Sciences, The University of Georgia, Athens, GA, 30602, USA; Food Animal Health Research Program, Ohio Agricultural Research and Development Center, The Ohio State University, Wooster, OH 44691, USA

**Keywords:** *Clostridium perfringens*, Vaccine, Nanoparticle, Immunity, Poultry, Necrotic enteritis

## Abstract

Currently, there is no commercial vaccine to control *Clostridium perfringens* (CP) or necrotic enteritis – a clinically and economically devastating disease. Two chitosan-nanoparticle encapsulated CP (CNP-CP) vaccines using extracellular proteins (ECP) and cell wall proteins (CWP) were synthesized: a CNP-CP ECP + CWP vaccine (SC vaccine) and a CNP-CP ECP + CWP surface-adsorbed CWP vaccine (SCC vaccine). The experiment comprised a complete randomized design of 3 treatments replicated 5 times: SC, SCC and nonimmunized control. Broilers orally gavaged with SC or SCC vaccine were primed and boosted with 70μg dose at 3- and 14-days post-hatch (dph). SDS-PAGE analysis revealed bands at 54.7 and 84.7 kDa in the ECP and 17 bands for CWP fraction. There were no differences in body weight gain, feed conversion ratio and mortality rate between treatments. At 17dph, serum of birds in the SC and SCC groups had higher neutralizing antibodies (*P*<0.01) compared to the control group. At 17dph, there was an increase in bile anti-ECP IgA levels in the SC vaccinated birds and a non-significant increase in SCC vaccinated birds compared to control. At 17dph, bile specific anti-CP CWP IgA levels were greater (*P*<*0.05*) in both immunized groups compared to control. At 17 and 21dph, serum anti-ECP and anti-CP CWP IgY levels in SC and SCC birds were comparable to the control. At 21dph, CD4+/CD8+ T-cell ratio in SC and SCC vaccinated groups were elevated (*P* ≤ *0.01*) compared to control. At 17dph, SC and SCC vaccinated birds had a significant reduction *(P* ≤ *0.001)* in α-toxin levels in caecal contents compared to control. Caecal α-toxin levels remained reduced at 21dph *(P* < *0.05)* in SC birds and numerically reduced in SCC vaccinated birds compared to control. Jejunal CP load in SCC birds was significantly reduced by 1.4 Log_10_ copy numbers of CP/g compared to control and no differences were observed in liver CP load between immunized and non-immunized birds. SC and SCC immunization did not alter TNF-α, IL-10, iNOS, IL-6 or IL-17 mRNA expression. At 17 and 21dph, SC and SCC immunized birds demonstrated greater sera bactericidal activity compared to control. CNP-SC and CNP-SCC immunization induced specific immune response against *C. perfringens* and reduced CP colonization via oral route of administration.

## INTRODUCTION

*Clostridium perfringens* (CP) is an anaerobic Gram positive, spore-forming, non-motile, rod-shaped bacterium found in diversified environments. Recently updated, CP strains are toxinotyped from A to G based on encoding combination of major toxins they produce; namely, α-toxin, β-toxin, ε-toxin and ι-toxin, enterotoxin (CPE) and NetB [1]. Necrotic enteritis (NE) is an enteric bacterial disease of poultry primarily caused by CP. Accompanying high morbidity and mortality of $US 6 billion in losses a year [2] prepares NE as a significant clinical and economically devastating disease of the poultry industry. In-feed antimicrobial growth promoters (AGPs) have well-controlled NE, however due to rising concerns of antimicrobial resistance, AGPs have been withdrawn from majority of the world [3]. Therefore, an alternative control measure to NE is needed. Presently, no commercial vaccine exists to control CP or NE.

The host-pathogen interaction of CP is complex as several predisposing factors contribute to the spontaneous outbreak of the disease. The majority of the 23 identified toxins produced by CP are encoded by large plasmids that range in size from ~45 kb to ~140 kb [4, 5]. These plasmid-encoded toxins are often closely associated with mobile elements. It has been reported that a CP strain can bear up to three different toxin plasmids, with a single plasmid carrying up to three distinct toxin genes [6]. Importantly, many toxin plasmids are closely related, suggesting a common evolutionary origin [7, 8]. In addition, CP secretes several growth inhibiting factors and hydrolytic proteins that are potentially involved in the pathogenesis of NE [9]. The supernatant of CP contains immunoreactive proteins such as alpha-toxin, glyceraldehyde-3-phosphate dehydrogenase, pyruvate:ferredoxin oxidoreductase and fructose 1, 6-biphosphate aldolase [9]. As such, CP supernatant proteins have been investigated as potential vaccine antigens to prevent clinical and subclinical NE [10–12]. It is also well acknowledged that many proteins are specifically anchored in the Gram-positive cell wall. Gram-positive bacteria have evolved numerous mechanisms by which they can immobilize proteins on their surface [13–16]. The functional importance of the Gram-positive cell wall proteins in distinct cellular compartments and bacteria-host interaction has been emphasized by several trials [15, 16–18] and appear to be significant in mediating pathogen interaction.

Research on CP vaccines have focused on parenteral killed vaccines or orally administered live attenuated strains. Oral inactivated vaccination would be an eminent value to the poultry industry due to the several disadvantages accompanying live and parenteral vaccines [19]. More importantly, oral vaccination mimics the route of CP infection. However, the proteolytic environment of the gastrointestinal tract and inefficient antigen presentation and processing in the gut results in the failure of orally delivered inactivated vaccines.

Biodegradable nanoparticles provide an innovative strategy for vaccine delivery to mucosal sites in terms of their dual function, both as adjuvants and delivery vehicles. Nanoparticles facilitate antigen uptake by professional antigen-presenting cells (APCs), maintain slow and controlled antigen release, protect vaccine’s antigen(s) from enzymatic degradation, and potentiate adaptative immunity [20]. Chitosan, α(1–4)2-amino 2-deoxy β-D glucan, is a natural mucoadhesive polysaccharide derived from partial deacetylation of chitin present in abundance in the shells of crustaceans [21, 22]. Chitosan has previously reported to stimulate mucosal secretory IgA, IgG, tumor necrosis factor-α (TNF-α), interlukin-6 (IL-6) and interferon-γ (IFN-γ) production [21, 22]. In addition, chitosan has shown to induce toll-like receptor (TLR) 2 or 5-mediated innate responses, rendering chitosan attractive carriers of novel mucosal vaccine antigens [23]. Chitosan nanoparticles are targeted delivery vehicles that can protect the antigen cargo, increase the antigen gut residence time, and decrease the gastric and intestinal degradation of the antigens [22, 23].

In this study, two chitosan nanoparticle *C. perfringens* (CNP-CP) vaccines containing native extracellular protein (ECP) and cell wall proteins (CWP) were synthesized: (1) an encapsulated ECP and CWP vaccine (SC) and (2) an encapsulated ECP and CWP surface-adsorbed CWP vaccine (SCC). The long-term goal is to develop a safe biodegradable NP-based oral CP vaccine for the poultry industry that can effectively reduce CP pathogenic load and NE severity in broilers. It is hypothesized that the oral CNP-CP subunit vaccines will elicit an immune response in broilers.

## MATERIALS AND METHODS

### Clostridium perfringens strain

*Clostridium perfringens* toxinotype A necrotic enteritis-induced field strain *CP6* was obtained from Dr. M. França, the laboratory of The Poultry Diagnostic Research Centre (PDRC) of The University of Georgia, Athens, GA, USA.

### Isolation and Identification and culture conditions of Clostridium perfringens

*C. perfringens* necrotic enteritis-induced field strain *CP6* was inoculated into blood agar base (Sigma-Aldrich®, MO) supplemented with 5% of sheep blood (LAMPIRE biological laboratories, PA) and incubated anaerobically with CO_2_ anaerobic pack (AnaeroPack® System, Mitsubishi Gas Chemical Company, INC., NY) in sterile bags at 37°C for 24 hours. Isolated CP colonies (1-2 colonies) were sub-cultured in 50 mL tubes containing thioglycolate broth with resazurine (Sigma-Aldrich®, MO) supplemented with 2% beef extract (Bioworld, bioPLUS Inc.) and incubated at 37 ºC for 24h. A second subculture of 10% v/v of culture with thioglycolate broth was performed for 24 h. At 24 h of subculture, a total of 200 mL subculture was further inoculated into 2 L of thioglycolate broth for 24 h at 37 °C under anaerobic conditions. At 24 h of incubation, the optical density of the culture, measured at 600nm with a spectrophotometer (BioTek Epoch spectrophotometer, Scarborough, ME; GEN5 3.03 software) was 0.6. The supernatant was collected after centrifugation at 11,420 ×*g* for 30min and filtered using a 0.22 μm filter (Sarstedt, Newton, NC).

### Native extracellular protein purification of C. perfringens field strain CP6

Supernatant proteins in the native form were precipitated from the supernatant using 80% w/v (NH_4_)_2_SO_4_ (Amresco, OH) at 4 °C overnight. The native supernatant was concentrated and purified by a process modified from the method by [24]. The native supernatant was reconstituted in 40 mL of PBS, concentrated and desalted 5X using a 10 kDa Amicon® 8400 Ultra filter membrane (Sigma Aldrich, St. Louis, MO). The purified and dialysed native CP extracellular protein was stored at −20 °C until further use.

### Native cell wall protein purification of C. perfringens field Strain CP6

Cell wall proteins fraction was prepared by the method reported earlier for another Gram-positive bacterium [25] with minor modifications. Briefly, *C. perfringens* cells were grown on thioglycolate broth at 37°C and 20 mL of 24h culture harvested in the exponential growth phase (OD_600 nm_=0.600) by centrifugation at 2,100×*g* for 10 min. The cells were washed with 62.5 mM Tris–HCl (pH 6.8) buffer to remove traces of extracellular proteins, then the bacteria were pipetted up-and-down for 3 min in 1 ml of the same buffer supplemented with 0.1% (v/v) Triton X-100 in order to solubilize the proteins associated to the cell wall. The suspension was centrifuged at 20,200×*g* for 5 min at 4°C. The supernatant was filtered through a 0.22 μm syringe filter (Milipore, India). Native cell wall proteins were precipitated from the supernatant using 80% w/v (NH_4_)_2_SO_4_ (Amresco, OH) at 4 °C overnight. The supernatant was reconstituted in 40 mL of PBS, concentrated and desalted 5X using a 10 kDa Amicon® 8400 Ultra filter membrane (Sigma Aldrich, St. Louis, MO). The supernatant was collected as the cell wall protein fraction and stored −20 °C until further use.

### C. perfringens ECP and CWP quantification

Chitosan-nanoparticle *C. perfringens* ECP and CWP were collected by centrifugation at 27,000× *g* for 20 minutes and were lyophilized with 5% sucrose as a cryoprotectant and stored at −80 °C until further use. A final concentration of 39 mg/mL and 10 mg/mL of *C. perfringens* CP6 native ECP and native CWP, respectively was obtained.

### Preparation of chitosan nano-encapsulated C. perfringens ECP +CWP vaccine (CNP-CP SC) and chitosan nano-encapsulated C. perfringens ECP+CWP surface-absorbed CWP (CNP-CP SCC)

Loaded chitosan nanoparticles were prepared in collaboration with the Food Animal Health Research Program, at The Ohio State University, USA. The CNP-CP SC and CNP-CP SCC were prepared by ionic gelation as described earlier [26]. *C. perfringens* ECP and CWP (2.5 mg each) were reconstituted in 1 mL of 1X MOPS buffer. The reconstituted ECP and CWP were added dropwise to 5 mL of a solution of 10 mg/mL of chitosan polymer (Sigma Aldrich, St. Louis, MO) in nano-pure water under magnetic stirring. A 12.5 mg of sodium tripolyphosphate (TPP) dissolved in a 25 mL 1X PBS, was added to the above mixture dropwise for chitosan: TPP ratio of 4:1. *C. perfringens* CWP were added to the above mixture dropwise to synthesize the CNP-CP SCC nanoparticle by surface-adsorbing the CWP. The SC and SCC nanoparticles were collected by centrifugation at 10,000×g for 30 min, lyophilized and stored at −20 °C until further use.

### Vaccination of birds with CNP-CP SC and CNP-CP SCC vaccines

The experimental procedure was approved by the Institutional Animal Care and Use Committee (IACUC), University of Georgia and was completed at The University of Georgia Poultry Science Research Complex over a period of 21days effective the date of chick arrival. The trial comprised a total of 45 male Cobb 500-day hatched broilers, raised in battery cages in an environmentally controlled poultry house. The experimental day-hatched chicks originated from the same breeding flock and hatchery. Birds were raised under standard managemental practices, the facility included a closed-system poultry house with poultry cages, air fans exhaust ventilators and cooling pads with a continuous lighting program provided. The experiment constituted a complete randomized design of 3 treatments replicated 5 times. Birds were equally distributed into 3 treatments: control (non-immunized), chitosan-nanoparticle encapsulated *C. perfringens* ECP and CWP vaccine (SC) and chitosan-nanoparticle encapsulated *C. perfringens* ECP and CWP surface-absorbed CWP (SCC). Birds in the SC and SCC groups were orally gavaged with a total of 70 μg of CNP-CP-SC and CNP-CP-SCC, respectively at 3 and 14 days of age post-hatch (dph). Starter and grower corn-soybean meal-based diets were formulated to meet the nutrient requirements for the modern Cobb 500 (Cobb-Vantress, Inc.). Birds welfare and health status were monitored daily and feed and water were offered *adlibitum* for the duration of the trial. At 17 and 21 days-post hatch, one bird per pen was individually weighted, sacrificed in accordance with IACUC and necropsy was performed.

### Production performance of birds vaccinated with CNP-CP SC and CNP-CP SCC vaccines. vaccines

Body weight and feed intake were measured at 0, 7, 14, and 21 dph, and body weight gain (BWG) and feed conversion ratio (FCR) were calculated.

### SDS-PAGE of C. perfringens ECP and CWP used in vaccine synthesis

*C. perfringens* ECP and CWP used to prepare the vaccine were separated on a 12 % SDS-PAGE gel run at 100V for 1 hour and 30 min visualized with Coomassie R250 staining. The loaded samples comprised of 2 μg, 4 μg and 8 μg protein. Band visualization was obtained via Gel DocTM XR+ and molecular weight (kDA) analyses of bands was achieved via BioRad Image Lab 6.0.1 after lane profiling and adjustment and comparison to molecular weight standard (Prometheus Prestained Protein Ladder, Genesse Scientific)

### Ex-vivo recall-response of splenic PBMCs of birds vaccinated with CNP-CP SC and CNP-CP SCC vaccines

Spleen samples were collected from one bird/cage on 17 and 21 dph (n=5). Single-cell suspensions of peripheral blood mononuclear cells (PBMCs) were collected and plated at 5 × 10^5^ cells per well in triplicates per sample in 100 μL of RPMI-1640 (Sigma Aldrich, St. Louis, MO) supplemented with 10% fetal bovine serum and 1% Penicillin and Streptomycin. *C. perfringens* ECP of 0.05 mg/mL, 0.1 mg/mL, 0.25 mg/mL or 0.5 mg/mL was added to each well (in duplicates) and incubated for 5 d at 37 °C in the presence of 5% CO_2_. Splenocytes stimulated with 0.0 mg/mL of antigen served as positive control and wells with no splenocytes served as negative control. PBMC proliferation was measured using MTT tetrazolium 3-(4,5-dimethylthiazol-2-yl)-2,5-diphenyltetrazolium bromide) assay as previously described [27]. Optical density was measured at OD_570_ using a spectrophotometer as described above. Percentage of PBMC viability was calculated using the following formula [28]: % Viability = [(Absorbance_exp_-mean (Absorbance_neg-control_) / mean (Absorbance_pos-control_)] × 100.

### Anti-C. perfringens ECP neutralizing antibodies in serum of birds vaccinated with CNP-CP SC and CNP-CP SCC vaccines

Splenocytes were used to assess *C. perfringens* toxin neutralization of serum antibodies in immunized birds. Seeded splenocytes in 96-well plates at 100 μl/well in RPMI-1640 supplemented with 10% fetal bovine serum and 1% Penicillin and Streptomycin at 1×10^6^ splenocytes/well in triplicates per sample were incubated at 37°C, 5% CO_2_ for 6 hours. Non-adhered cells were discarded. Sera samples from of 5 birds/treatment collected at 17 and 21 dph were subjected to 1:20 dilution in RPMI-1640 media used to seed cells. A neutralization plate containing 75 μL of 1 mg/mL *C. perfringens* ECP was incubated with either 75 μL of 1:20 dilution of serum, or RPMI-1640 media only serving as positive control. A set of wells with 150 μL of RPMI-1640 media only was maintained as negative control. Incubation was maintained for 1 h at 37 °C, with rotating agitation to neutralize toxins. A 100 μL of the neutralizing solution was added to splenocytes and further incubated for 2 d at 37 °C and 5% CO_2_. Splenocyte proliferation was measured using the MTT tetrazolium (3-(4,5-dimethylthiazol-2-yl)-2,5-diphenyltetrazolium bromide) assay as described above and absorbance values were reported as OD_570nM_.

### Anti-CP ECP and anti-CP CWP specific IgY and IgA antibody levels in serum and bile of birds vaccinated with CNP-CP SC and CNP-CP SCC vaccines

Serum and bile were collected from one bird per cage (n=5) at 17 and 21 dph. Anti-CP ECP and anti-CP CWP specific IgY and IgA antibody levels in serum and bile were determined by enzyme-linked immunosorbent assay (ELISA). ELISAs were developed and optimized by checkerboard titration. Briefly, *C. perfringens* native ECP or native CWP were coated on ELISA plates (Nunc MaxisorpTM, ThermoFisher Scientific, Waltham, MA) at 10 μg/mL (IgA) or 25 μg/ mL (IgY) diluted in 0.02 M sodium carbonate-bicarbonate coating buffer, pH 9.6 and incubated overnight at 4°C. Bile was diluted to 1:200 and serum was diluted to 1:20 in PBS containing 2.5% non-fat dry milk, and 0.5% Tween 20 (VWR, Radnor, PA) and samples were incubated for 1 hour. Horseradish peroxidase (HRP) conjugated polyclonal goat anti-chicken IgG (Bethyl, Montgomery, TX) at 1:20,000 dilution or HRP-conjugated polyclonal goat anti-chicken IgA (SouthernBiotech, Birmingham, AL) at 1:10,000 was used as a secondary antibody and plates were incubated for 1 h, then washed with PBS containing 0.5% Tween 20 (PBST) and aspirated. A 3,3’,5,5’-tetramethylbenzidine (TMB) substrate was added to each well and incubated for 15 min. A 2M sulfuric acid was added to stop the reaction after 10 min. Optical density (OD) was measured as absorbance at 450nM using a spectrophotometer (Biochek, Scarborough, ME) and values are reported as OD_450_.

### Alpha-toxin load in ceca and C. perfringens load in jejunum and liver of birds vaccinated with CNP-CP SC and CNP-CP SCC vaccines

Caecal content, jejunum and liver samples (n=5) were collected from one bird/cage at 17 and 21 dph and stored at −80°C until DNA extraction. Caecal contents, jejunum and liver samples (approximately 100 mg) were washed 3 times with 1× PBS. The cell pellet was resuspended in EDTA and treated with 20 mg/mL lysozyme for 45 min at 37°C, followed by treatment with lysis buffer containing 20% SDS and 0.1 mg/mL proteinase K (Sigma Aldrich, St Louis, MO) for 10 min at 80°C. The samples were incubated with 5 μL of RNase at 37°C for 30 min. The cell lysate was incubated with 6 M sodium chloride on ice for 5 min. The supernatant was collected after centrifugation at 400× g for 10 min. The DNA in the supernatant was precipitated with isopropanol and washed once in ice-cold ethanol. The DNA pellet was resuspended in TE buffer (10 mM Tris-HCl, 1 mM EDTA, pH 8.0) and stored at −20°C until further use. The concentration of the isolated DNA was determined by using NanoDrop™ 2000c Spectrophotometer (ThermoFisher Scientific, Waltham, MA, USA). The DNA samples were diluted to a final concentration of 100 ng/μl. The *C. perfringens* alpha-toxin (CPA) load in the caeca was analyzed by real-time PCR, using primers 5’-GCAGCAAAGGTAACTCTAGCTAACT-3’ and 3’-CCTGGGTTGTCCATTTCCCATT-5’) with an annealing temperature was 55°C. The Ct was determined by iQ5 software (Bio-Rad, Hercules, CA, USA) when the fluorescence rises exponentially 2-fold above background. Alpha-toxin Ct values were expressed as mean Ct values subtracted from 40 cycles and presented as 40-Ct. A standard curve was drawn with amplicon DNA to determine the copy number for *C. perfringens* and the amplicon DNA of *C. perfringens* was quantified in an Epoch Microplate Spectrophotometer (BioTec Instruments, Winooski, VT) and were serially diluted in deionized water to obtain a concentration of 1 ng, 0.1 ng, 0.01 ng, 1 pg, 0.1 pg, and 0.01 pg [48]. The serially diluted amplicon was analyzed by real-time PCR for *C. perfringens* primer. The PCR efficiency and the slope and intercept of the standard curve was determined by the IQ5 software. The PCR efficiency of the *C perfringens* standard curve analysis was 97.7%. The slope and intercept of the standard curve for was Cq= −3.418 × Log (DNA) +12.679. The quantity of the DNA amplicon in caecal contents was calculated using the standard curve after determining the Ct value by real-time PCR. The *C. perfringens* load in jejunum and liver was analyzed by real-time PCR, using primers 5’-AAAGGAAGATTAATACCGCATAA-3’ and 5’-ATCTTGCGACCGTACTCCCC-3’) with an annealing temperature was 60°C. The Ct was determined by iQ5 software (Bio-Rad, Hercules, CA, USA) when the fluorescence rises exponentially 2-fold above background. The copy number of *C. perfringens* was analyzed as described earlier [29] with minor modification. The concentration of *C. perfringens* DNA was converted to an amplicon equivalent for calculation of copy numbers. The mean mass of the DNA amplicon was calculated using the following equation described earlier [30]. Mean mass = (number of base pairs × 607.4 + 157.9). The number of base pairs in *C. perfringens* primers DNA amplicon was 172. The copy numbers were calculated using the formula as described earlier [29]: Copy number = DNA quantity/ (mean mass × 6.023 × 10^23^). *C. perfringen*s load in caecal contents were presented as Log_10_ copy numbers/g.

### TNF-α, IL-10, iNOS, IL-6 and IL-17 mRNA levels in caecal tonsils of birds vaccinated with CNP-CP SC and CNP-CP SCC vaccines

Caecal tonsil samples were collected at 17 and 21 dph (n=5). Total RNA was extracted by using TRIzol reagent (Invitrogen, Carlsbad, CA, USA). The isolated RNA was dissolved in Tris-EDTA (pH 7.5) buffer, and the concentration was determined by using NanoDrop™ 2000c Spectrophotometer (ThermoFisher Scientific, Waltham, MA, USA). The cDNA synthesis was achieved with 2 μg of total RNA. The mRNA transcripts analyzed for the relative expression of TNF-α, IL-10, iNOS, IL-6 and IL-17 with housekeeping gene *β-*actin mRNA by qPCR using the iQ™ SYBR® Green Supermix (Bio-Rad, Hercules, CA, USA). Forward and reverse primers and PCR conditions for RT-PCR are listed in Table 1. The threshold cycle (Ct) values were determined by iQ5 software (Bio-Rad, Hercules, CA, USA). The fold change from the reference was calculated using the 2^(Ct Sample—Housekeeping)^/2^(Ct Reference—Housekeeping)^ comparative Ct method, where Ct is the threshold cycle [31]. The Ct was determined by iQ5 software (Bio-Rad, Hercules, CA, USA) when the fluorescence rises exponentially 2-fold above the background.

**Table 1.**
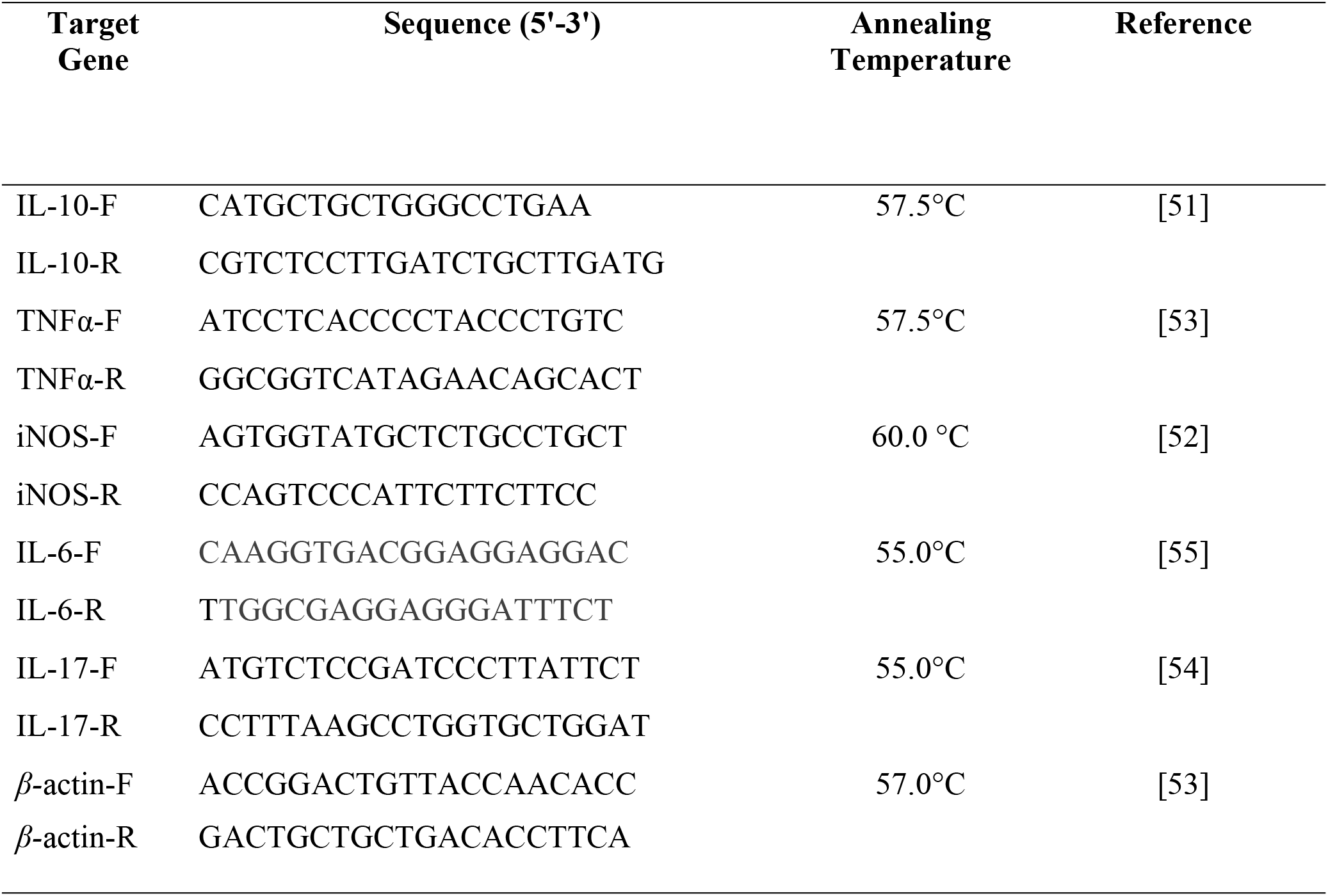
Primers and PCR conditions for qRT-PCR.

### CD4^+^, CD8^+^, and CD4^+^/CD8^+^ T-cell ratio in caecal tonsils of birds vaccinated with CNP-CP SC and CNP-CP SCC vaccines

Caecal tonsil samples were teased over a 0.4 μm cell strainer (Sigma–Aldrich, St. Louis, MO) with approximately 2 mL RPMI-1640 to obtain a single-cell suspension. Single-cell suspensions of caecal tonsils from vaccinated and non-vaccinated birds at 17 and 21 dph were concentrated for lymphocytes by density centrifugation over Histopaque (1.077 g/mL; Sigma–Aldrich, St. Louis, MO). For CD4^+^/CD8^+^ T-cell ratio analysis, single-cell suspensions of the cecal tonsils (1 × 10^6^ cells) were incubated with PE-conjugated mouse anti-chicken CD4, FITC-conjugated mouse anti-chicken CD8 (Southern Biotech, Birmingham, AL) at 1:200 dilution, and unlabeled mouse IgG at 1:200 dilution in a 96-well plate for 20 min. After incubation, cells were washed twice by centrifugation at 400 × g for 5 min using wash buffer (1× PBS, 2 mM EDTA, 1.5% FBS) to remove unbound primary antibodies. After washing, cells were analyzed using cytosoft software (Guava Easycyte, Millipore, Billerica, MA). The CD4^+^, CD8^+^ and CD4^+^/CD8^+^ T-cell ratio was analyzed after gating cells based on forward-scatter and side-scatter plot for lymphocytes.

### Serum bactericidal activity of birds vaccinated with CNP-CP SC and CNP-CP SCC vaccines

To assess CNP-CP ECP and CWP vaccines’-induced antibody function in conjunction with complement in sera, a serum bactericidal assay was performed. Briefly, sera samples from each treatment group were pooled and diluted in a 1: 8 in thioglycolate broth supplemented with 2% beef extract, then incubated at 37°C with 24h *C. perfringens CP6* bacterial culture grown in thioglycolate broth supplemented with 2% beef extract (Log_10_ 6.67/ml; OD_600_=0.9513) for 1 hour in a 1:1 dilution. A 40 ul of incubated sera-CP culture mixture was inoculated on sheep blood agar supplemented with 2% beef extract (in triplicates) for Log_10_ CFU/ml enumeration of CP.

### Statistical Analysis

Statistical differences were determined using one-way ANOVA followed by Tukey’s HSD test for mean separation. Statistical analyzes were performed using proper procedures of JMP Pro 15 (SAS Institute Inc., USA; 2019) with *P-values* <*0.05* considered statistically significant.

## RESULTS

### Production performance of birds vaccinated with CNP-CP SC and CNP-CP SCC vaccines. vaccines

There were no significant differences (*P* > 0.05) in the mean BWG, FI, FCR and mortality between any of the treatment groups at 7, 14 and 21 dph (Table 2).

**Table 2.** Perfromance parameters of male Cobb 500 broilers vaccinated with SC and SCC vaccines.

|  | *0-7 dph* |  |  | *7-14 dph* |  |  | *14-21 dph* |  |  |  |
| --- | --- | --- | --- | --- | --- | --- | --- | --- | --- | --- |
| Treatment | BWG | FI | FCR | BWG | FI | FCR | BWG | FI | FCR | Mortality |
|  | (g) | (g) |  | (g) | (g) |  | (g) | (g) |  | (%) |
| **Control** | 264 | 550 | 2.01 | 574 | 914 | 1.61 | 704 | 1052 | 1.54 | 0 |
| **SC** | 282 | 549 | 1.97 | 516 | 937 | 1.85 | 670 | 1050 | 1.63 | 0 |
| **SCC** | 250 | 534 | 2.26 | 551 | 1019 | 1.85 | 800 | 1175 | 1.49 | 0 |
| **SEM** | 17.6 | 27.0 | 0.23 | 40.9 | 58.3 | 0.12 | 93.3 | 95.8 | 0.10 | - |

Broilers in SC and SCC groups were vaccinated at 3 and 14 dph with 70 μg of chitosan-encapsulated *C. perfringens* extracellular + cell wall proteins vaccine and chitosan-encapsulated extracellular + cell wall proteins with surface-adsorbed cell wall proteins vaccine, respectively. Broilers in control group were non-immunized. Dph: Days post-hatch; BWG: Body weight gain; FI: Feed intake; FCR: Feed conversion ratio; SEM: Standard error of mean.

### SDS-PAGE of C. perfringens ECP and CWP used in vaccine synthesis

SDS-PAGE of 24h incubated broth ECP and CWP from *C. perfringens* (CP6) associated field necrotic enteritis strain is illustrated in S1 Fig. *C. perfringens* Type A cultured in thioglycolate broth enriched with 2% beef extract demonstrated bands for ECP (in lane 2-4; at 2 μg/ml, 4 μg/ml and 8 μg/ml, respectively) and CWP (in lane 5-7; at 2 μg/ml, 4 μg/ml and 8 μg/ml, respectively) post extraction using 80% w/v (NH_4_)_2_SO_4_ and purification. Bands for ECP fraction of this strain had a prominent band at 54.7kDa and 84.7 kDa in lanes 2-4. Lane 5-7 illustrate 17 bands for CWP fraction for CP6 (S1 Fig).

### Ex-vivo recall-response of splenic PBMCs of birds vaccinated with CNP-CP SC and CNP-CP SCC vaccines

Primary splenocytes were pulsed with *CP6* ECP to determine the recall response at 17 and 21 dph. PBMCs from birds in the SC and SCC group at 17 dph had a significant increase (*P<0.05)* in T-lymphocyte proliferation at 0.10 mg/mL ECP stimulation compared to control group. PBMCs from birds in the SCC group at 17 d dph had a significant increase (*P<0.05)* in T-lymphocyte proliferation at 0.25 and 0.50 mg/mL ECP stimulation compared to SC and control group (Fig. 1).

**Fig. 1.**
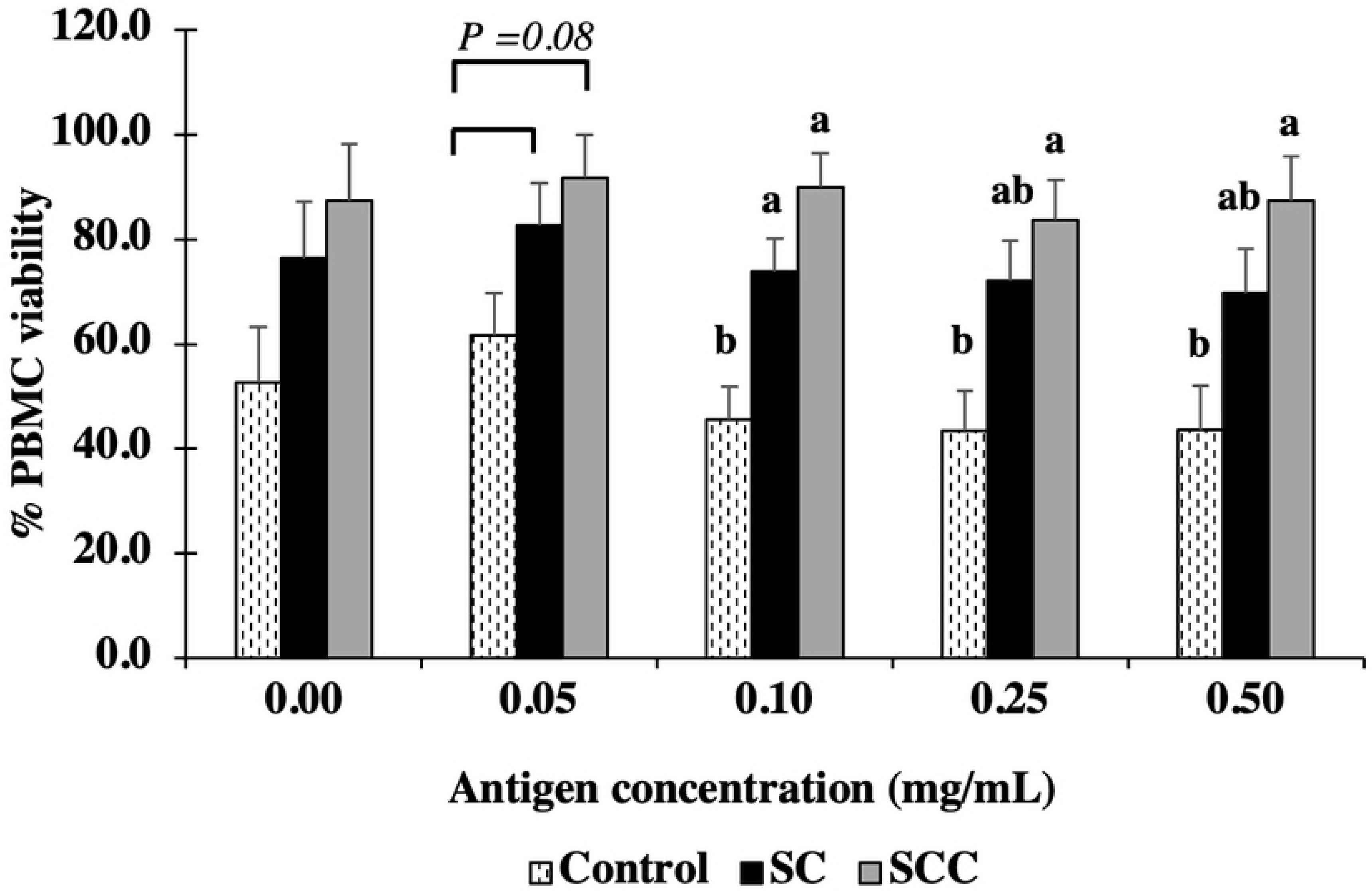
Ex-vivo recall-response of splenic PBMCs of chickens vaccinated with CNP-CP SC and CNP-CP SCC vaccines. Broilers in SC and SCC groups were vaccinated at 3 and 14 dph with 70 μg of chitosan-encapsulated *C. perfringens* extracellular + cell wall proteins vaccine and chitosan-encapsulated extracellular + cell wall proteins with surface-adsorbed cell wall proteins vaccine, respectively. Broilers in control group were non-immunized. Splenocytes PBMCs of broilers at 17 dph were harvested and stimulated with 0.05, 0.1, 0.25 and 0.5 mg/mL of C. perfringens extracellular proteins for 72 h. Recall response was measured using MTT at OD_570nM_ to report percentage of PBMC viability. Broilers in control group were non-immunized. Mean+SEM. ^a,b^Means with differing alphabetical superscript are significantly different (*P*<*0.05*).

### Anti-C. perfringens ECP neutralizing antibodies in serum of birds vaccinated with CNP-CP SC and CNP-CP SCC vaccines

The level of anti-CP ECP neutralizing antibodies in the serum was measured by the ability of the serum to reverse the CP-induced decrease in splenocyte proliferation by incubating the serum with CP ECP. At 17 dph, splenocyte incubated with ECP and serum from SC and SCC groups had higher proliferation compared to splenocyte incubated with only ECP and compared to splenocytes incubated with ECP and serum from the control group (*P* < 0.01) At 21 dph, splenocytes incubated with ECP and serum from SCC group had higher proliferation compared to splenocytes incubated with only ECP and compared to splenocytes incubated with ECP and serum from the control group or serum from SC group (*P* < 0.01) (Fig. 2).

**Fig. 2.**
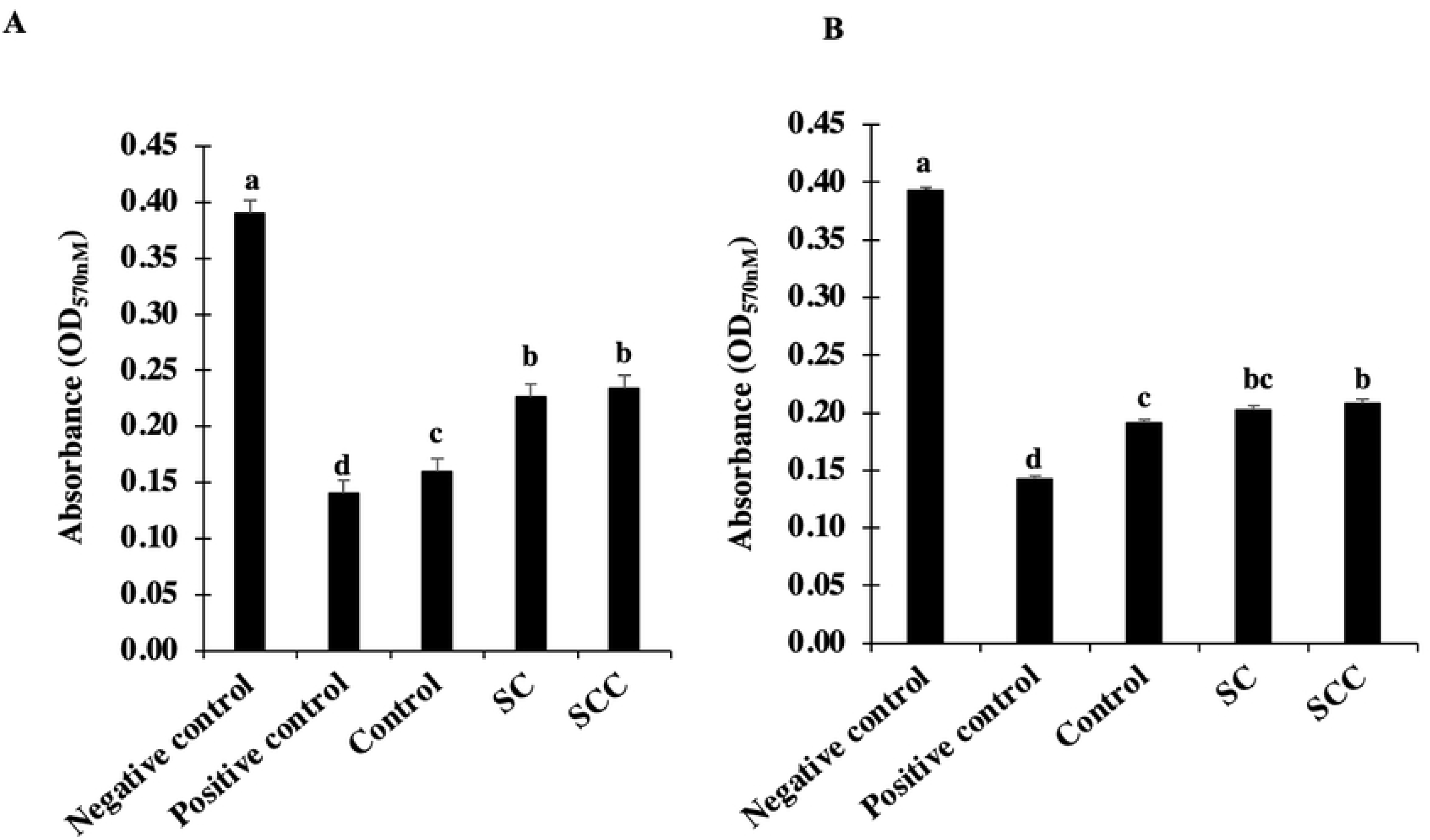
Anti-C. perfringens ECP neutralizing antibodies in serum of chickens vaccinated with CNP-CP SC and CNP-CP SCC vaccines. Broilers in SC and SCC groups were vaccinated at 3 and 14 dph with 70 μg of chitosan-encapsulated *C. perfringens* extracellular + cell wall proteins vaccine and chitosan-encapsulated extracellular + cell wall proteins with surface-adsorbed cell wall proteins vaccine, respectively. Broilers in control group were non-immunized. Serum from 5 birds/treatment were collected at (A) 17 and (B) 21 dph and were incubated with splenocytes and native ECP from *C. perfringens CP6*. Optical density (OD) values were measured at 570mM. Mean+SEM. ^a,b,c^Means with differing alphabetical superscript are significantly different (*P*<*0.001*).

### Anti-CP ECP and anti-CP CWP specific IgY and IgA antibody levels in serum and bile of birds vaccinated with CNP-CP SC and CNP-CP SCC vaccines

At 17 and 21 dph, broilers in the SC and SCC groups had comparable sera anti-CP ECP IgY and anti-CP CWP IgY to that in control group (Fig. 3A, 3B). Bile anti-CP ECP IgA in SC immunized broilers were significantly greater than those in control birds at 17 dph, but not at 21 dph (Fig. 3C). Bile anti-CP CWP IgA in SC and SCC immunized groups were significantly greater than those in control birds at 17 dph, and not at 21 dph (Fig. 3D).

**Fig. 3.**
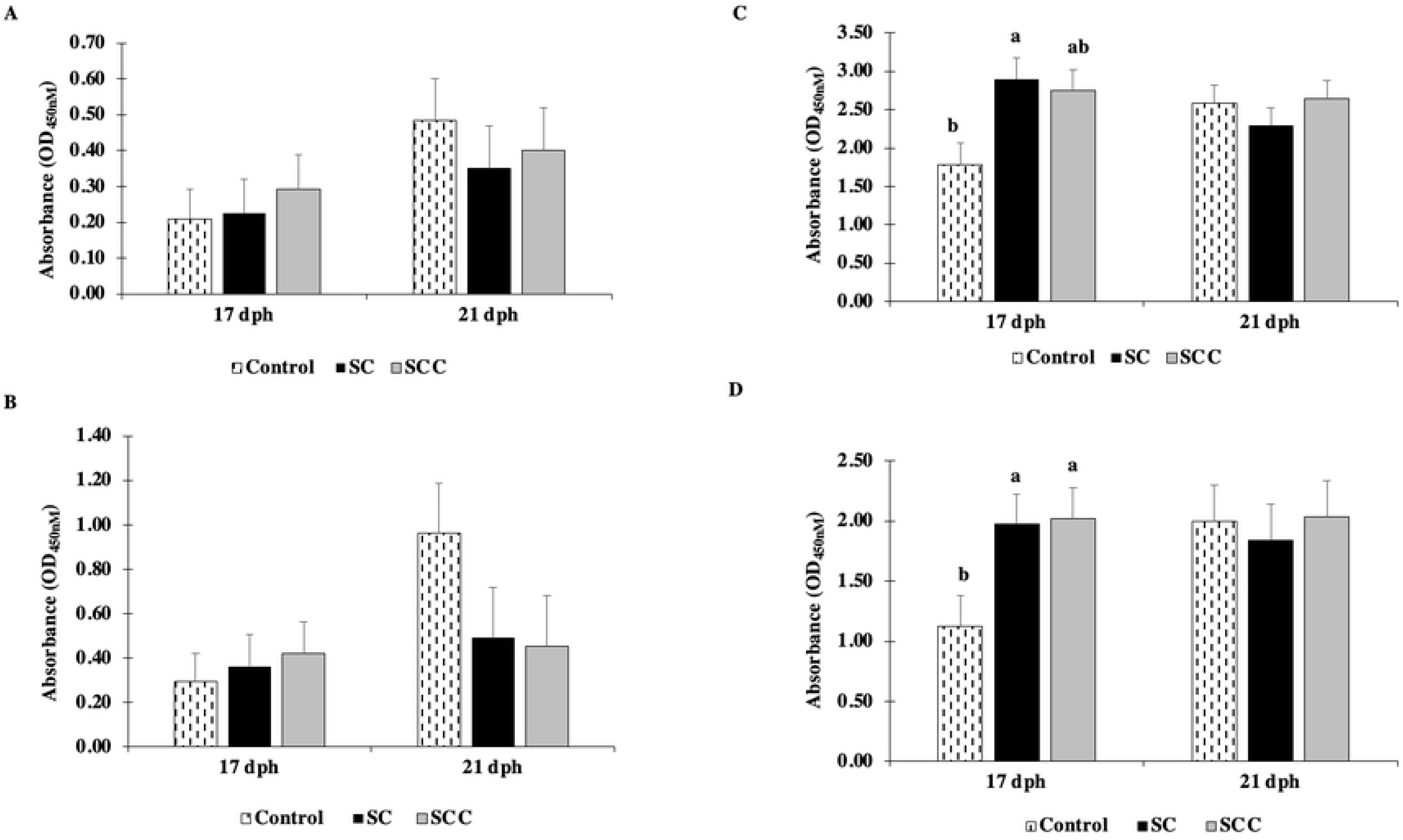
Anti-C. perfringens ECP and anti-C. perfringens CWP specific IgY and IgA antibodies of birds vaccinated with CNP-CP SC and CNP-CP SCC vaccine. Broilers in SC and SCC groups were vaccinated at 3 and 14 dph with 70 μg of chitosan-encapsulated *C. perfringens* extracellular + cell wall proteins vaccine and chitosan-encapsulated extracellular + cell wall proteins with surface-adsorbed cell wall proteins vaccine, respectively. Broilers in control group were non-immunized. Sera samples (n=5) were collected to analyze for (A) anti-ECP IgY and (B) anti-CWP IgY at 17 and 21 dph. Bile samples (n=5) were collected to analyze for (C) anti-ECP IgA and (D) anti-CWP IgA at 17 and 21 dph. Optical density values (OD) were measured at 450nM. Means ± SEM. ^a,b^Means with differing alphabetical superscript are significantly different (*P* < *0.05*).

### C. perfringens alpha-toxin load in ceca of birds vaccinated with CNP-CP SC and CNP-CP SCC vaccines

At 17 dph, birds immunized with SC and SCC vaccines had significant reduction (*P* ≤ *0.001)* in alpha-toxin load compared to those in control group. By 21 dph, SC immunized birds maintained a significant reduction (*P* < *0.05)* in alpha-toxin levels in cecal contents, while SCC birds had a non-significant reduction in alpha-toxin compared to birds that were non-immunized (Fig. 4).

**Fig. 4.**
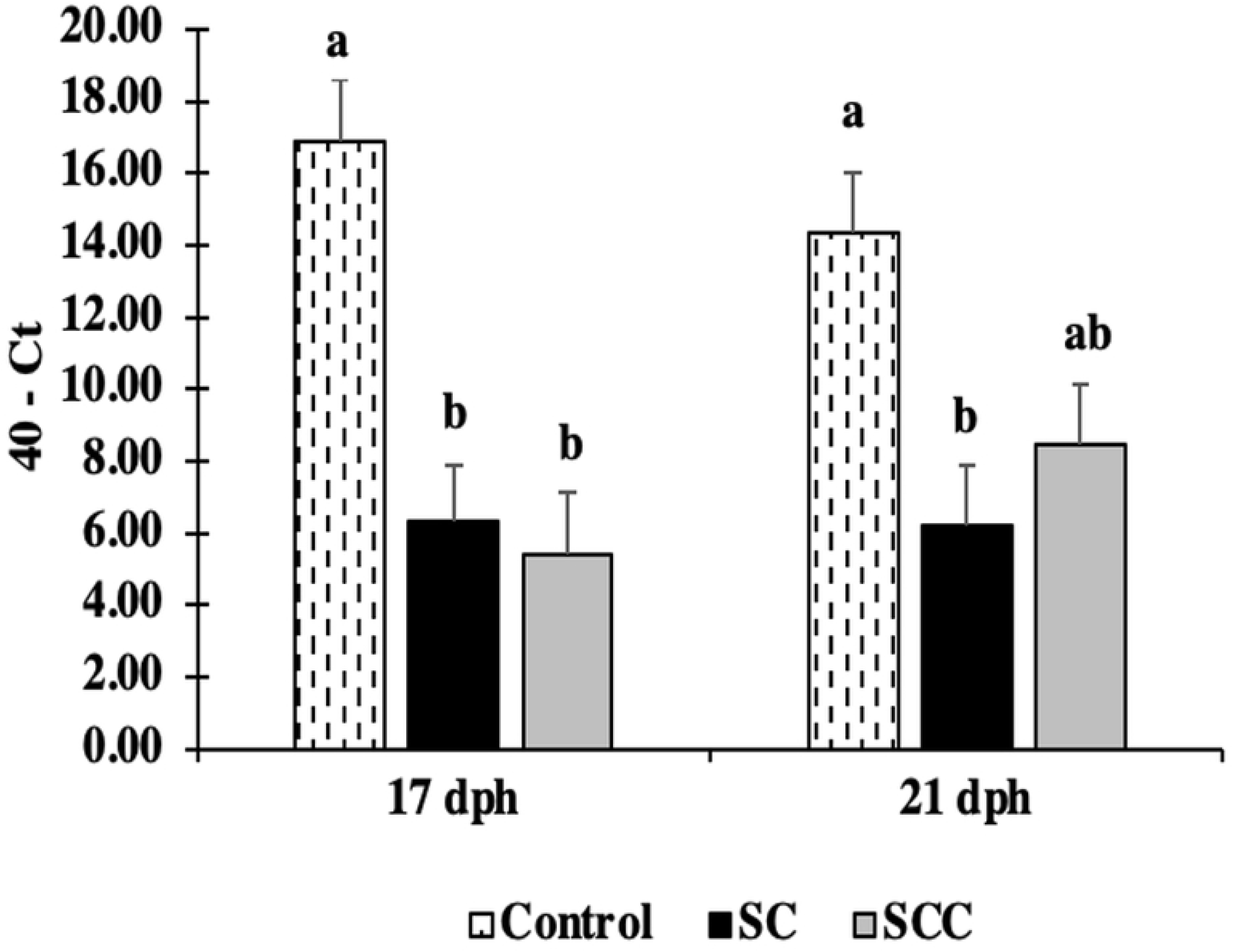
C. perfringens alpha-toxin load in caeca at 17 and 21 dph of birds vaccinated with CNP-CP SC and CNP-CP SCC vaccines. Broilers in SC and SCC groups were vaccinated at 3 and 14 dph with 70 μg of chitosan-encapsulated *C. perfringens* extracellular + cell wall proteins vaccine and chitosan- encapsulated extracellular + cell wall proteins with surface-adsorbed cell wall proteins vaccine, respectively. Broilers in control group were non-immunized. Alpha-toxin mean cycle threshold (Ct) values (n=5) were subtracted from total of 40 qPCR cycles. Means ± SEM. ^a,b^Means with differing alphabetical superscript are significantly different (*P* < *0.05).*

### C. perfringens load in jejunum and liver of birds vaccinated with CNP-CP SC and CNP-CP SCC vaccines

At 17 dph, birds immunized with SCC vaccines had significant reduction (*P* < *0.05)* in *C. perfringens* load in jejunum compared to those in SC and control group. By 21 dph, birds in SC and SCC had comparable levels of CP in jejunum compared to control (Fig. 5A). At 17 and 21 dph, levels of CP in liver of birds in SC and SCC groups were comparable to those in control (Fig. 5B).

**Fig. 5.**
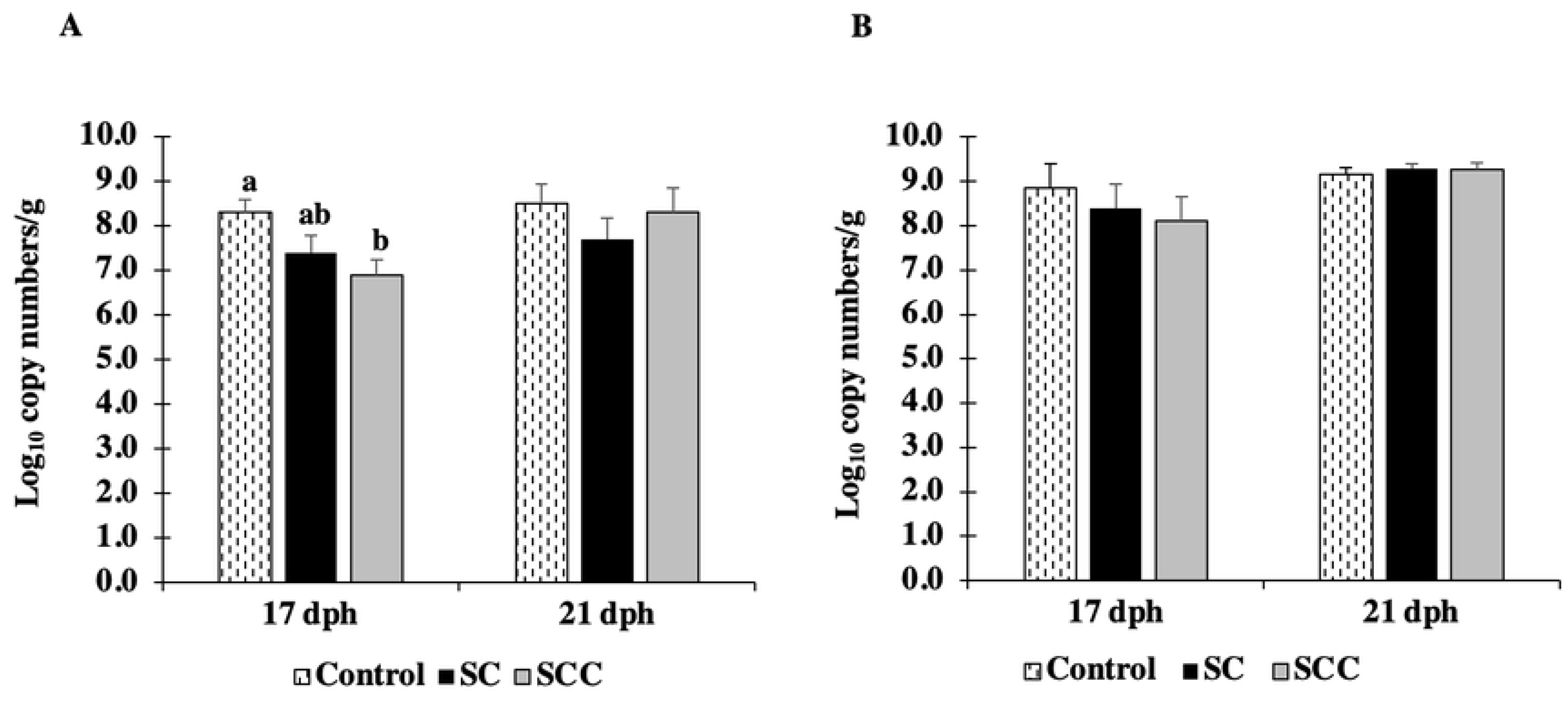
C. perfringens load in jejunum and liver of birds vaccinated with CNP-CP SC and CNP-CP SCC vaccines. Broilers in SC and SCC groups were vaccinated at 3 and 14 dph with 70 μg of chitosan-encapsulated *C. perfringens* extracellular + cell wall proteins vaccine and chitosan-encapsulated extracellular + cell wall proteins with surface-adsorbed cell wall proteins vaccine, respectively. Broilers in control group were non-immunized. *C. perfringens* load was represented as Log_10_ copy numbers/g in (A) jejunum and (B) liver at 17 and 21dph (n=5) via q-PCR. Means+SEM. ^a,b^Means with differing alphabetical superscript are significantly different (*P*<*0.05*).

### TNF-α, IL-10, iNOS, IL-6 and IL-17 mRNA expression in caecal tonsils of birds vaccinated with CNP-CP SC and CNP-CP SCC vaccines

At 17 and 21dph, caecal tonsils of birds from the SC and SCC group showed no significant difference in mRNA expression of TNF-α, IL-10, iNOS compared to those in the control group (*P* >0.05). A non-significant upregulation in IL-6 at 17 dph and IL-17 at 21 dph is observed in SCC group compared to the control group (*P* >0.05) (Fig. 6A, 6B, 6C and 6D).

**Fig. 6.**
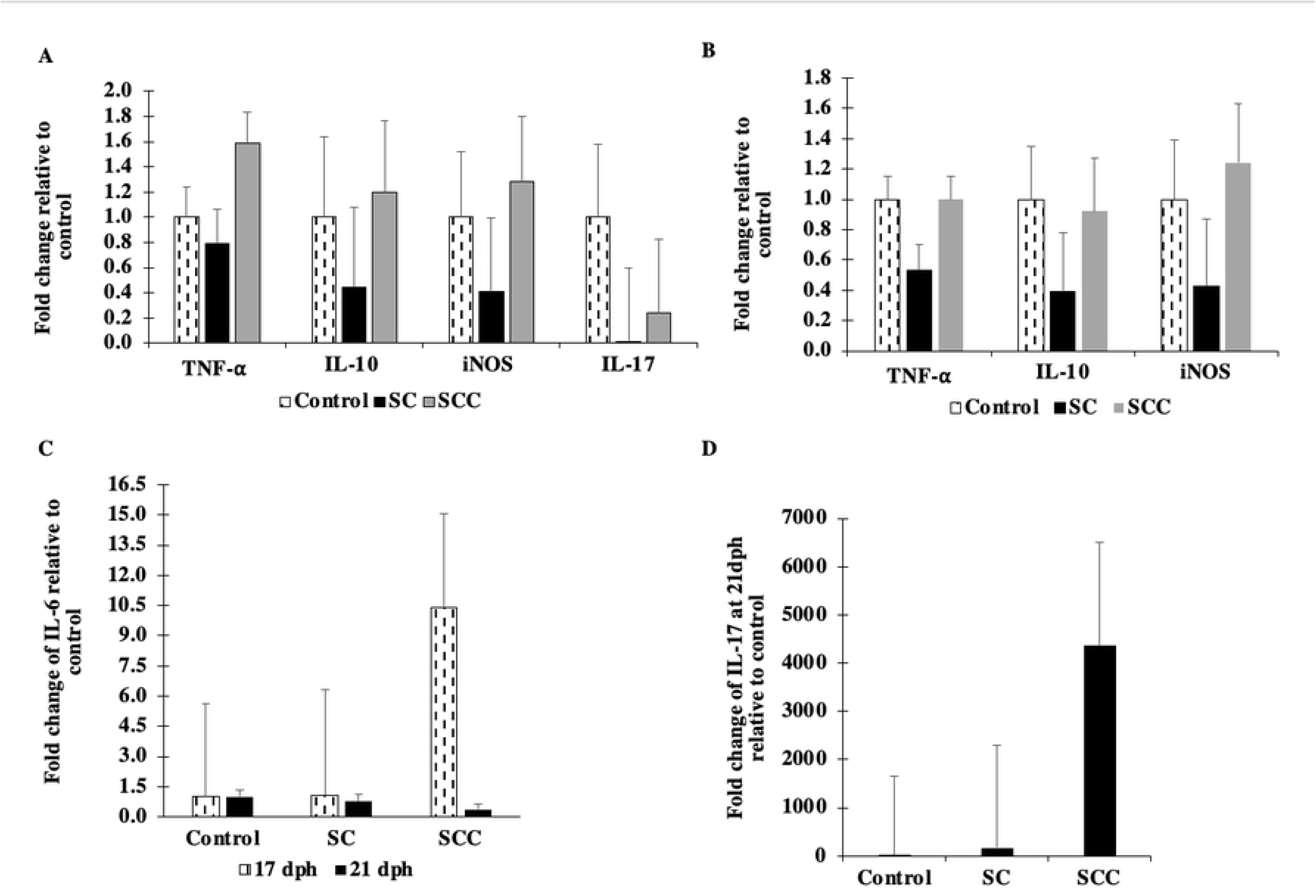
TNF-α, IL-10, iNOS, IL-6 and Il-17 mRNA levels in caecal tonsils of birds vaccinated of birds vaccinated with CNP-CP SC and CNP-CP SCC vaccines. Broilers in SC and SCC groups were vaccinated at 3 and 14 dph with 70 μg of chitosan-encapsulated *C. perfringens* extracellular + cell wall proteins vaccine and chitosan- encapsulated extracellular + cell wall proteins with surface-adsorbed cell wall proteins vaccine, respectively. Broilers in control group were non-immunized. TNF-α, IL-10 and iNOS, IL-6 and IL-17 mRNA levels in caecal tonsils (n=5) analyzed by qRT-PCR. Means+SEM. (A) TNF-α, IL-10, iNOS and IL-17 at 17 dph; (B) TNF-α, IL-10 and iNOS at 21 dph; (C) IL-6 at 17 and 21 dph; (D) IL-17 at 21dph.

### CD4^+^, CD8^+^ and CD4^+^/CD8^+^ T-cell ratio in caecal tonsils of birds vaccinated with CNP-CP SC and CNP-CP SCC vaccines

Caecal tonsils of birds immunized with *SC and SCC vaccine had a greater* CD4^+^/CD8^+^ T-cell ratio population approaching significance at 17 dph *(P=0.06)* compared to those in the control group. By 21 dph, SC and SCC immunized birds had greater (*P* ≤ *0.01*) population of CD4^+^/CD8^+^ in caecal tonsils compared to broilers in the control group (Fig. 7A, 7B and 7C).

**Fig. 7.**
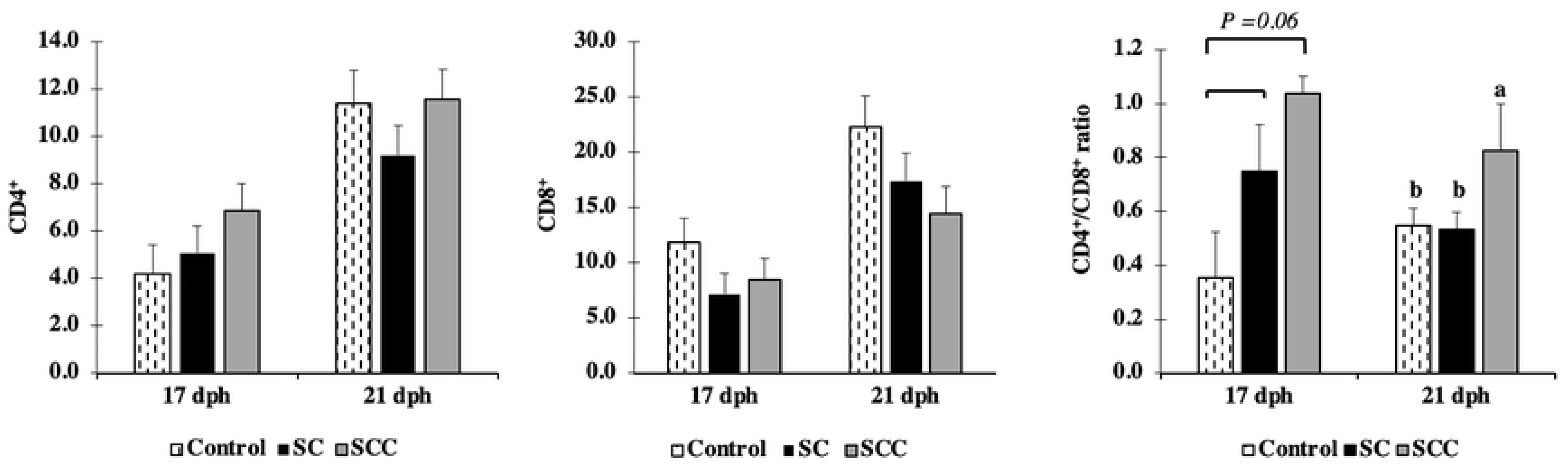
CD4^+^, CD8^+^ and CD4^+^/CD8^+^ T-cell ratio in caecal tonsils at 17 and 21 dph of birds vaccinated with CNP-CP SC and CNP-CP SCC vaccines. Broilers in SC and SCC groups were vaccinated at 3 and 14 dph with 70 μg of chitosan-encapsulated *C. perfringens* extracellular + cell wall proteins vaccine and chitosan- encapsulated extracellular + cell wall proteins with surface-adsorbed cell wall proteins vaccine, respectively. Broilers in control group were non-immunized. CD4^+^, CD8^+^, CD4^+^/CD8^+^ ratio in caecal tonsils were analyzed by flow cytometry after gating cells based on forward-scatter and side-scatter plot for lymphocytes. Means ± SEM. a,bMeans with differing alphabetical superscript are significantly different (*P* ≤ *0.01*).

### Serum bactericidal activity of birds vaccinated with CNP-CP SC and CNP-CP SCC vaccines

At 17 dph, sera of birds immunized with SC and SCC vaccines had a greater bactericidal activity (*P* < *0.05*) compared to those in control group (Fig. 8). Sera of SC immunized birds incubated with Log_10_ 6.67 CFU/ml CP demonstrated no detectable CFUs of CP at 17dph, while sera of SCC immunized birds incubated with Log_10_ 6.67 CFU/ml CP had reduced CP by 1.6 Log_10_ CFU/ml compared to control of 1.2 Log_10_ CFU/ml reduction. Similarly, at 21 dph, sera of birds in SC and SCC groups had a greater bactericidal activity (*P* < *0.05*) compared to non-immunized birds (Fig. 8). Sera of SC immunized birds incubated with Log_10_ 6.67 CFU/ml CP demonstrated 1.60 Log_10_ CFU/ml reduction in CP at 21 dph compared to control of 0.9Log_10_ CFU/ml reduction, while sera of SCC immunized birds incubated with Log_10_ 6.67 CFU/ml CP demonstrated no detectable CFUs at 21 dph.

**Fig. 8.**
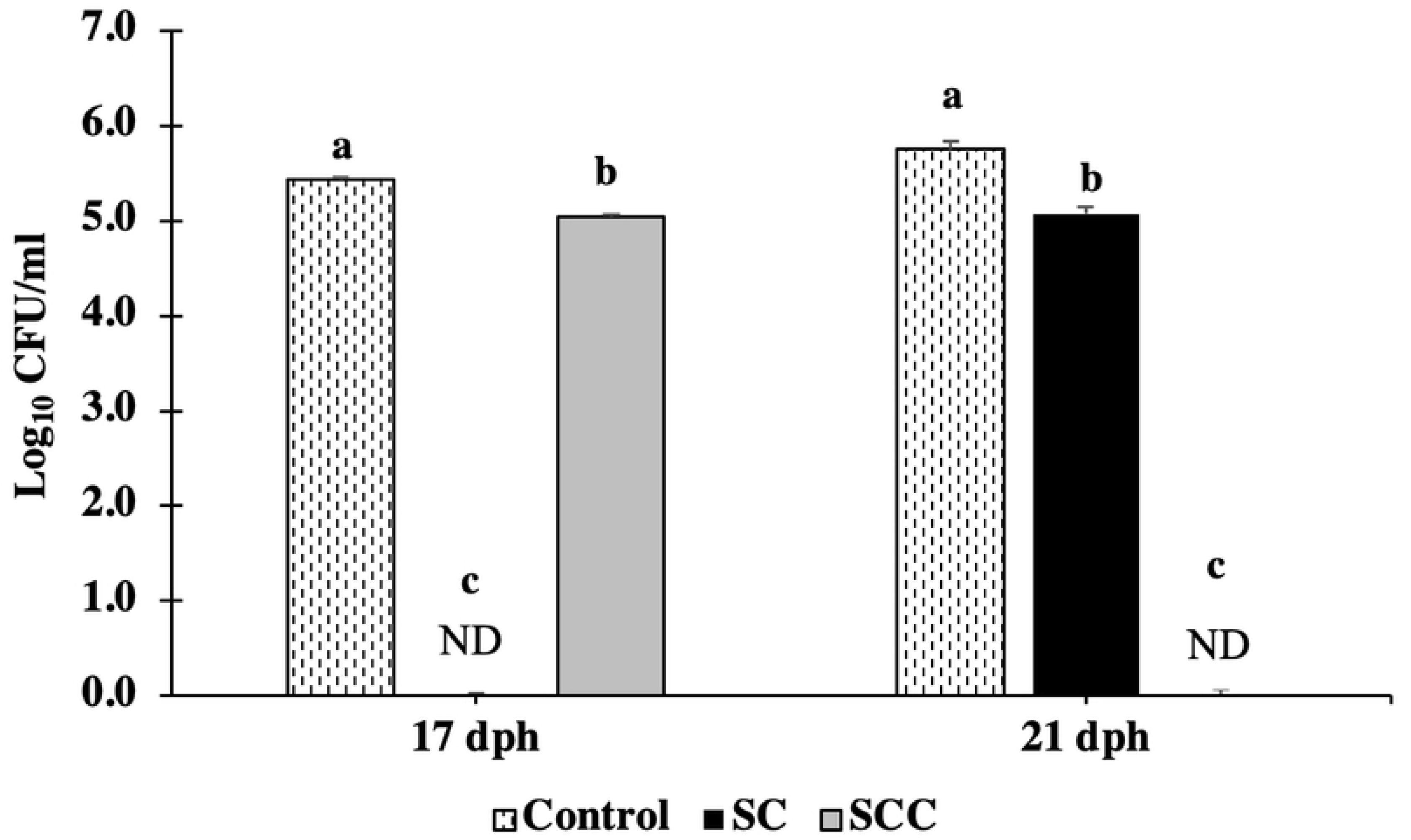
Serum bactericidal activity at 17 and 21 dph of birds vaccinated with CNP-CP SC and CNP-CP SCC. Broilers in SC and SCC groups were vaccinated at 3 and 14 dph with 70 μg of chitosan-encapsulated *C. perfringens* extracellular + cell wall proteins vaccine and chitosan-encapsulated extracellular + cell wall proteins with surface-adsorbed cell wall proteins vaccine, respectively. Broilers in control group were non-immunized. Sera samples at 17 and 21 dph from each treatment (n=5) were pooled and incubated with broth culture *C. perfringens* (Log_10_6.67/ml; OD_600_=0.9513) at a dilution of 1:1 for 1 hour. Data represented as mean Log_10_ colony forming units (CFU)/mL on sheep blood agar supplemented with 2% beef extract (technical replicates = 3). Means+SEM. ^a,b,c^Means with differing alphabetical superscript are significantly different (*P* < *0.01)*.

## DISCUSSION

Presently, no commercial vaccine exists for CP or NE. In this study, two oral chitosan nanoparticles *C. perfringens* vaccines encapsulating ECP and cell wall subunits were synthesized: a CNP-CP ECP+CWP vaccine and a CNP-CP ECP +CWP surface-adsorbed CWP vaccine. Virulence properties of different CP strains are determined by their capability of secreting various proteinaceous toxins and enzymes; causing different forms of tissue damage. The *C. perfringens* isolate, CP6, used for this study has been isolated from field NE case and was shown by qPCR to be α-toxin positive and *NetB* negative (data not shown). Earlier studies have described *NetB* negative isolates recovered from chickens with NE [32].

The CNP-CP vaccines in this study were synthesized using CP native ECP and CWP subunits. Antigens selected for subunit immunization ideally include properties of extracytoplasmic localization, safety and immunogenicity, and conservation among different pathogenic strains [16]. *Clostridium perfringens* ECP and CWP are immunogenic and conserved between several strains of CP [9, 16]. In addition, several CP ECP and CWP are extracytoplasmic [16, 33]. Extracytoplasmic proteins comprise variety of functions including nutrient uptake, adhesion, chemosensing, envelop biogenesis, extracellular degrading enzymes and cell-surface virulence factors [34]. The purified native ECP of CP6 fraction used in vaccine synthesis included a prominent band at 54.7 kDa and 84.7 kDa, while the purified CWP fraction of this strain constituted 17 different bands.

This experiment was carried to investigate the efficacy and immunogenicity of *in-vivo* orally administered CNP-CP ECP and CWP vaccines in broilers. In the present study, vaccination of broilers with CNP-CP SC and CNP-CP SCC vaccines increased anti-CP ECP neutralizing antibodies in the serum, reduced alpha-toxin caecal load and CP load in jejunum.

The FI, BWG, FCR were not significantly different between immunized and non-immunized birds. Similarly, the mortality rate was not different between treatment groups throughout the 21-day experimental period, indicating that both SC and SCC candidate vaccines were safe with no adverse effects on bird performance. Earlier studies reported that birds orally inoculated CNP did not have any effect on mortality [26, 27].

The vaccine-induced recall response to ECP antigen was reflected by PBMC viability. Splenocyte PBMCs of SC and SCC vaccinated birds demonstrated greater percentage of PBMC viability compared to the control group at 0.10 mg/mL ECP stimulation. While, splenocyte PBMCs of SCC vaccinated birds had significantly greater percentage of PBMC viability at 0.25 and 0.50 mg/mL than SC vaccinated birds and those in the control group. Such antigen-specific induced recall response after mucosal vaccination is in accordance with previous studies of PBMC responses against CP antigen stimulation [35] and may be due to differences in epitope density [36] or antigen processing and presentation, and/or dependent on antigen type and stimulated concentration.

In the current study, neutralizing anti-specific ECP antibodies in sera of SC and SCC groups were significantly higher at 17dph compared to control. Fukata et al. [37] reported that neutralization of alpha toxin in purified toxin or *C. perfringens* culture supernatant with a hyperimmune serum against alpha toxin prevents the induction of NE. The current observations are in agreement with our previous studies [35, 38] where birds vaccinated with polyanhydride or chitosan CP ECP vaccine had higher ECP-neutralizing antibodies in sera compared to those in control group. Although birds in the SC and SCC groups had no differences in the total anti-CP ECP-specific IgY or anti-CP CWP-specific IgY from those in control group, the anti-CP ECP-specific IgA levels of birds immunized with SC vaccine were significantly greater at 17 dph with a coefficient of variation (CV) of 7.58% compared to control group (CV=53.78%). Birds immunized with SCC had non-significant increase of anti-CP ECP-specific IgA with a CV of 13.6% compared to control birds. Good and poor uniformity are CV of <20 or >20%, respectively, indicating homogeneity or heterogeneity of vaccination [49]. In addition, the anti-CP CWP-specific IgA levels of birds immunized with SC or SCC vaccine were greater at 17 dph with CV of 18.6 and 28.1%, respectively compared to those in the control group (CV=56.6%). Mirsha and Smyth [39] reported that oral live CP immunization of broilers induced serum and mucosal antibody responses at 22 days of age.

The CD4^+^/CD8^+^ T-cell ratio in caecal tonsils of SC and SCC vaccinated birds approached significance (*P=0.06*) at 17 dph with greater population than those in control birds. By 21 dph, CD4^+^/CD8^+^ ratio in SC and SCC groups were significantly increased (*P* ≤ *0.01*) compared to nonimmunized birds. This reflects a cell-mediated immune response induced by the SC and SCC vaccines. According to Wang et al. [40], CD4^+^/CD8^+^ ratio in lamina propria lymphocytes from ileum were significantly greater in broilers challenged with subclinical NE (SNE) supplemented Lactobacillus *johnsonii* BS15 than those in control challenged SNE. Combination of carvacrol, cinnamaldehyde, and capsicum oleoresin was shown to stimulate cell-mediated immunity by CD4^+^, CD8^+^ T-cells that enhanced immunity against coccidiosis by stimulating the innate and humoral immune responses [41]. Johnston et al. [42] reported that CD4^−/−^ and pIgR^−/−^ mice challenged with *C. difficile* were protected from *C. difficile* infection, while MHCII^−/−^ mice were not protected. Anti-toxin IgA antibodies were found to be protective in *C. difficile* infected CD4^−/−^ mice, which lack IgG class switching. In contrast, MHCII−/− mice; which are unable to generate IgG or IgA class switching, were not protected against *C. difficile* infection; suggesting IgG or IgA is required, yet not both. The greater PBMC viability in this study mounting a sufficient immune response to CP antigen may be attributed to a T-cell dependency reflected by the greater CD4^+^/CD8^+^ T-cell ratio and the anti-specific CP IgA antibodies observed in SC and SCC immunized birds. Together, these findings suggest that T-cell mediated response may be needed in order to generate a mucosal anti-CP specific antibody response.

To investigate the effect of SC and SCC vaccines on *C. perfringens* load in broilers, qPCR enumeration of α-toxin in cecal contents and CP in jejunum and liver was performed. At 17 dph, immunized SC and SCC birds demonstrated a significant reduction *(P* ≤ *0.001)* in α-toxin in caecal contents compared to those nonimmunized. The α-toxin load remained reduced *(P* < *0.05)* at 21 dph in SC immunized birds and numerically reduced in SCC vaccinated birds compared to the control group. While, birds immunized with SCC vaccine had a reduction of 1.4 Log_10_ copy numbers of CP/g of jejunum at 17 dph compared to birds in the control group. Alpha toxin is involved in the assimilation of nutrients by degrading membrane phospholipids. The production of alpha toxin is regulated by the *luxS* gene through the common quorum-sensing signal autoinducer 2 [43], indicating that alpha toxin is involved in competitive exclusion of other enteric bacteria and in intestinal colonization. Thus, the reduction in alpha-toxin could cause interruption of these functions and inhibit CP growth.

Reduction in CP intestinal load through vaccination may prevent the incidence of NE. The study observed that CNP-CP ECP + CWP subunit vaccination increased sera anti-ECP neutralizing antibodies. Anti-ECP neutralizing antibodies are expected to decrease intestinal CP proliferation. In addition, sera of birds immunized with SC and SCC vaccines had a greater sera bactericidal activity towards CP broth culture compared to those in control group. Zekarias et al. [44] reported alpha toxin-neutralizing serum antibodies in broilers orally immunized with 10^9^ CFU recombinant attenuated *Salmonella*-based delivery of the C-terminal domain of alpha toxin. In the same study, sera of immunized broilers suppressed CP bacterial growth when added to CP broth cultures [44].

Bactericidal activity in serum appears to be of importance for protection against systemic infections [45–47, 50]. In addition, induction of mucosal IgA antibodies is associated with local protection against invasion by pathogenic bacteria through the mucosal membranes [40]. Immunization is an important strategy in minimizing enteric pathogens in poultry, mitigating morbidity and mortality rates and foodborne risk of human diseases. The ability of CNP-SC and CNP-SCC to reduce alpha-toxin and CP load in broilers’ intestines and elicit an enhanced immune response is a promising element of the candidate vaccines. Further studies are needed to assess the protective effects of CNP-CP ECP+CWP vaccines, particularly under field conditions.

## ACKNOWLEDGEMENT

The author would like to thank Dr. Monique França at the PDRC, Athens, GA for field *C. perfringens strain CP6.* The author would like to acknowledge Gabriel Akerele, Keila Acevedo, Mohamed Murtada, Theros Ng, Jarred Oxford and Ragini Reddyvari from The University of Georgia, Athens, GA for their help in sample collections.

## REFERENCES

[1] Rood, J. I. et al. (2018) Expansion of the *Clostridium perfringens* toxin-based typing scheme. Anaerobe. 10.1016/j.anaerobe.2018.04.011.

[2] Wade, B. and Keyburn, A. 2015. The true cost of necrotic enteritis. World Poult. 31. 16–17.

[3] Bennett PM. 1995. The spread of drug resistance. In: Baumberg S Young JPW Wellington EMH Saunders JR editors. Pop Gen in Bacteria. Cambridge:317–44.

[4] Kiu, R., Caim, S., Alexander, S., Pachori, P. & Hall, L. J. (2017). Probing Genomic Aspects of the Multi-Host Pathogen Clostridium perfringens Reveals Significant Pangenome Diversity, and a Diverse Array of Virulence Factors. Front Microbiol 8, 2485.

[5] Theoret, J. R., Uzal, F. A. & McClane, B. A. (2015). Identification and characterization of Clostridium perfringens beta toxin variants with differing trypsin sensitivity and in vitro cytotoxicity activity. Infect. Immun. 83, 1477–1486.

[6] Bannam T Yan X Harrison P Seemann T Keyburn A Stubenrauch C etal. 2011. Necrotic enteritis-derived Clostridium perfringens strain with three closely related independently conjugative toxin and antibiotic plasmids. mBio 2(5):e00190–11.

[7] Miyamoto K Li J Sayeed S Akimoto S McClane BA. 2008. Sequencing and diversity analyses reveal extensive similarities between some epsilon-toxin-encoding plasmids and the pCPF5603 Clostridium perfringens enterotoxin plasmid. J. Bacteriol. 190:7178–7188.

[8] Sayeed S Li J McClane BA. 2010. Characterization of virulence plasmid diversity among Clostridium perfringens type B isolates. Infect. Immun. 78:495–504.

[9] Kulkarni, R. V. R. Parreira, S. Sharif, and J. F. Prescott. (2007). Immunization of Broiler Chickens against *Clostridium perfringens*-Induced Necrotic Enteritis. Clinical and Vaccine Immunology, p. 1070–1077.

[10] Crouch CF Withanage GS de Haas V Etore F Francis MJ. Safety and efficacy of a maternal vaccine for the passive protection of broiler chicks against necrotic enteritis. Avian Pathol 2010;39(6):489–97.

[11] Saleh N Fathalla SI Nabil R Mosaad AA. 2011. Clinicopathological and immunological studies on toxoids vaccine as a successful alternative in controlling clostridial infection in broilers. Anaerobe;17(6):426–30.

[12] Jiang Y Mo H Willingham C Wang S Park JY Kong W etal. Protection against necrotic enteritis in broiler chickens by regulated delayed lysis salmonella vaccines. Avian Dis 2015;59(4):475–85.

[13] Schneewind O Mihaylova-Petkov D Model P. Cell wall sorting signals in surface proteins of gram-positive bacteria. EMBO J. 1993;12(12):4803–4811.

[14] Berge A Kihlberg B M, Sjöholm A G, Björck L. Streptococcal protein H forms soluble complement-activating complexes with IgG, but inhibits complement activation by IgG-coated targets. J Biol Chem. 1997;272:20774–20781.

[15] Navarre WW Schneewind O. Surface proteins of gram-positive bacteria and mechanisms of their targeting to the cell wall envelope. Microbiol Mol Biol Rev. 1999;63(1):174–229.

[16] Alam SI Bansod S Kumar RB Sengupta N Singh L. (2009). Differential proteomic analysis of Clostridium perfringens ATCC13124; identification of dominant, surface and structure associated proteins. BMC Microbiol. 2009;9:162.

[17] Hiscox TJ Chakravorty A Choo JM etal. Regulation of virulence by the RevR response regulator in Clostridium perfringens. Infect Immun. 2011;79(6):2145–2153.

[18] Yang, H., Hoober, J. Regulation of *lip*A Gene Expression by Cell Surface Proteins in *Arthrobacter photogonimos*. Curr Microbiol 38, 92–95 (1999).

[19] Mot, D., Timbermont, L., Haesebrouck, F., Ducatelle, R., Van Immerseel, F., 2014. Progress and problems in vaccination against necrotic enteritis in broiler chickens. Avian Pathol. 43, 290–300.

[20] Lubben IM Verhoef JC Borchard G Junginger HE. (2001). Chitosan for mucosal vaccination. Adv Drug Deliv Rev 52:139–44.

[21] Read RC., Naylor SC., Potter CW., Bond J., Jabbal-Gill I., Fisher A., Illum L., Jennings R. Effective nasal influenza vaccine delivery using chitosan. Vaccine. 2005;23:4367–4374..

[22] Baaten B.J., Clarke B., Strong P., Hou S. Nasal mucosal administration of chitin microparticles boosts innate immunity against influenza a virus in the local pulmonary tissue. Vaccine. 2010;28:4130–4137.

[23] Bolhassani A., Javanzad S., Saleh T., Hashemi M., Aghasadeghi M.R., Sadat S.M. Polymeric nanoparticles: Potent vectors for vaccine delivery targeting cancer and infectious diseases. Hum. Vaccines Immunother. 2014;10:321–332. doi: 10.4161/hv.26796.

[24] Keyburn A.L., Boyce J.D., Vaz P., Bannam T.L., Ford M.E. et al. 2008. NetB, a new toxin that is associated with avian necrotic enteritis caused by *Clostridium perfringens*. PLoS Pathog. 2008;4(2) :e26.

[25] Tavares F. and Sellstedt A. DNase Activities of the Extracellular, Cell Wall-Associated, and Cytoplasmic Protein Fractions of Frankia Strain R43. Applied and Environmental Microbiology Nov 1997, 63 (11) 4597–4599.

[26] Dhakal, S., Renu, S., Ghimire, S., Shaan Lakshmanappa, Y., Hogshead, B.T., Feliciano-Ruiz, N., Lu, F. et al. 2018. Mucosal immunity and protective efficacy of intranasal inactivated influenza vaccine is improved by chitosan nanoparticle delivery in pigs. Frontiers Immunol. 9, 934.

[27] Zhao, K., Chen, G., Shi, X.M., Gao, T.T., Li, W., Zhao, Y., Zhang, F.Q., Wu, J., Cui, X., Wang, Y.F., 2012. Preparation and efficacy of a live newcastle disease virus vaccine encapsulated in chitosan nanoparticles. PLoS One 7 (12), 1–11.

[28] Kamiloglu, S, Sari, G, Ozdal T and Capanoglu E. (2020). Guidelines for cell viability assays. Food Front, 1;3.

[29] Cottyn, B., K. Heylen, J. Heyrman, K. et al. 2009. Pseudomonas cichorii as the causal agent of midrib rot, an emerging disease of greenhouse-grown butterhead lettuce in Flanders. Syst. Appl. Microbiol. 32:211–225.

[30] Shin, J. H., Y. Yue, and D. Duan. 2012. Recombinant adeno-associated viral vector production and purification. Methods Mol. Biol. 798:267–284.

[31] Schmittgen, T.D. and K.J. Livak, 2008. Analyzing real-time PCR data by the comparative CT method. Nature prot,3(6): 1101.

[32] Abildgaard, L., Sondergaard, T.T., Engberg et al. 2010. In vitro production of necrotic enteritis toxin B, NetB, by netB-positive and netB-negative Clostridium perfringens originating from healthy and diseased broiler chickens. Vet. Microbiol. 12.036.

[33] Sengupta, N, Syed Alam, S.I, Kumar B Kumar, R, Gautam, V, Kumar, S et al. (2010). Comparative Proteomic Analysis of Extracellular Proteins of Clostridium perfringens Type A and Type C Strains Infection and Immunity, 78 (9) 3957–3968.

[34] Wang G Chen H Xia Y Cui J Gu Z Song Y etal. (2013). How are the non-classically secreted bacterial proteins released into the extracellular milieu?. Curr Microbiol. 2013 Dec;67(6):688–95.

[35] Akerele G Ramadan N Renu S, Renukaradhya GJ Shanmugasundaram R Selvaraj RK. In vitro characterization and immunogenicity of chitosan nanoparticles loaded with native and inactivated extracellular proteins from a field strain of Clostridium perfringens associated with necrotic enteritis. Vet Immunol Immunopathol. 2020 Jun;224:110059.

[36] Wiertz E J H J, van Gaans-van den Brink J A M, Gausepohl H Procknicka-Chalufour A, Hoogerhout P Poolman J T. Identification of T cell epitopes occurring in a meningococcal class 1 outer membrane protein using overlapping peptides assembled with simultaneous multiple peptide synthesis. J Exp Med. 1992;176:79–88.

[37] Fukata, T., Y. Hadate, E. Baba, T. Uemura, and A. Arakawa. 1988. Influence of *Clostridium perfringens* and its toxin in germ-free chickens. Res. Vet. Sci. 44: 68–70.

[38] Ramadan, NR, Akerele G Renu S Renukaradhya GJ and Selvaraj RK. Oral poly(anhydride)-nanoparticle based *Clostridium perfringens* extracellular protein vaccine expressing *Salmonella* flagella reduces intestinal *Clostridium perfringens*. Forthcoming 2020.

[39] Mishra, N., Smyth, J.A., 2017. Oral vaccination of broiler chickens against necrotic enteritis using a non-virulent NetB positive strain of Clostridium perfringens type A. Vaccine 35 (49), 6858–6865.

[40] Wang H Ni X Qing X etal. Probiotic Enhanced Intestinal Immunity in Broilers against Subclinical Necrotic Enteritis. Front Immunol. 2017;8:1592.

[41] Naidoo V., L.J. McGaw, S.P.R. Bisschop, N. Duncan, J.N. Eloff. The value of plant extracts with antioxidant activity in attenuating coccidiosis in broiler chickens. Vet.Parasitol., 153 (2008), pp. 214–219.

[42] Johnston PF Gerding DN Knight KL. Protection from Clostridium difficile infection in CD4 T Cell- and polymeric immunoglobulin receptor-deficient mice. Infect Immun. 2014;82(2):522–531.

[43] Ohtani K Hayashi H Shimizu T. The luxS gene is involved in cell-cell signalling for toxin production in Clostridium perfringens. Mol Microbiol. 2002 Apr;44(1):171–9.

[44] Zekarias B Mo H Curtiss R 3rd. Recombinant attenuated Salmonella enterica serovar typhimurium expressing the carboxy-terminal domain of alpha toxin from Clostridium perfringens induces protective responses against necrotic enteritis in chickens. Clin Vaccine Immunol (5):805–16.

[45] Keo T Collins J Kunwar P Blaser M Iovine N (2011) Campylobacter capsule and lipooligosaccharide confer resistance to serum and cationic antimicrobials, Virulence, 2:1, 30–40

[46] Trebicka, E., Jacob, S., Pirzai, W., Hurley, B. P., & Cherayil, B. J. (2013). Role of antilipopolysaccharide antibodies in serum bactericidal activity against Salmonella enterica serovar Typhimurium in healthy adults and children in the United States. Clinical and vaccine immunology : CVI, *20*(10), 1491–1498.

[47] Sahin, O., N. Luo, S. Huang, and Q. Zhang. (2003). Effect of Campylobacter-specific maternal antibodies on Campylobacter jejuni colonization in young chickens. Appl. Environ. Microbiol. 69:5372–5379.

[48] Shanmugasundaram R Mortada M Murugesan GR Selvaraj RK. In vitro characterization and analysis of probiotic species in the chicken intestine by real-time polymerase chain reaction. Poult Sci. 2019 Nov 1;98(11):5840–5846.

[49] Quinn CP Semenova VA Elie CM Romero-Steiner S Greene C Li H etal. (2002). Specific, sensitive, and quantitative enzyme-linked immunosorbent assay for human Immunoglobulin G antibodies to anthrax protective antigen. Emerg. Infect. Dis. 8, 1103.

[50] Perkins BA Jonsdottir K Briem H Griffiths E Plikaytis BD Hoiby E etal. (1998). Immunogenicity of two efficacious outer membrane protein-based serogroup B meningococcal vaccines among young adults in Iceland. J Infect Dis.(3):683–91.

[51] Rothwell, L. Young, J. Zoorob, R., Whittaker, C., Hesketh, P. et al 2004. Cloning and characterization of chicken IL-10 and its role in the immune response to Eimeria maxima. J of Immuno;173(4): 2675–82.

[52] Selvaraj, R. Klasing K. C. Lutein and Eicosapentaenoic Acid Interact to Modify iNOS mRN Levels through the PPARγ/RXR Pathway in Chickens and HD11 Cell Lines Jour of Nutrition Vo136, 6.

[53] Markazi, A., Luoma A Shanmugasundaram R Mohnl M Raj Murugesan G, Selvaraj R. etal. 2018., Effects of drinking water synbiotic supplementation in laying hens challenged with *Salmonella*. Poul sci;. 97(10): p. 3510–3518.

[54] Min, W and Lillehoj H. 2012. Isolation and characterization of chicken inteleukin-17 cDNA. Journal of interferon & cytokine research 22:1123–1128

[55] Y.H. Hong, W. Song, S.H. Lee, H.S. Lillehoj, 2012. Differential gene expression profiles of β-defensins in the crop, intestine, and spleen using a necrotic enteritis model in 2 commercial broiler chicken lines, Poul Scie,91, 5,pp1081–1088,

